# Fusion of dysfunction muscle stem cells with myofibers induces sarcopenia in mice

**DOI:** 10.1101/2023.01.20.524967

**Authors:** Xun Wang, Prashant Mishra

## Abstract

Sarcopenia, or age-associated muscle atrophy, is a progressive condition which affects ∼10-30% of the human geriatric population (*1, 2*). A number of contributors to sarcopenia have been proposed, including the progressive loss of muscle stem cells (MuSCs) with age. However, studies in mice have provided evidence that MuSC depletion is not sufficient to induce sarcopenia (*3, 4*). We recently showed that in response to age-associated mitochondrial damage, MuSCs self-remove by fusing with neighboring myofibers, which depletes the stem cell population of damaged progenitors (*5*). Here, we show that MuSC-myofiber fusion is sufficient to initiate myofiber atrophy in mice, which limits their motor function and lifespan. Conversely, inhibition of MuSC-myofiber fusion blocks myofiber atrophy with age, with a concomitant increase in the maximum lifespan of animals. These findings suggest a model where the accumulation fusion of damaged MuSCs with adult myofibers is a key driving feature of sarcopenia, and resolves the findings that MuSC depletion on its own does not initiate myofiber atrophy.

---

In mammals, a number of studies indicate that MuSCs decline in numbers with age (*6-9*), suggesting that the limited regenerative capacity of aged muscle contributes to myofiber atrophy. However, in two recent studies (*3, 4*) in mice, MuSCs were depleted by conditional expression of a diphtheria toxin allele (DTA) at early age. Neither study observed an accelerated decline in myofiber size with age, calling into the question the contribution of MuSCs to age-associated atrophy.

In response to mitochondrial electron transport chain (ETC) dysfunction or environmental toxins, muscle stem cells are induced to directly fuse into neighboring adult myofibers (*5, 10*). These fusion events contribute to the decline in MuSC numbers with age; however, the consequences for the accepting myofiber are unknown (Fig. 1A). To address this, we used a conditional allele targeting Cox10, a critical assembly component of mitochondrial complex IV (*11*), combined with the tamoxifen-inducible Pax7-Cre^ERT2^ driver (*12*), to induce ETC dysfunction in adult MuSCs. Inhibition of complex IV function triggered the rapid removal of MuSCs, by inducing fusion with neighboring myofibers (Fig. S1A,B), consistent with previous results (*5*). We treated 6-week old mice with tamoxifen to induce Cox10 deletion in adult MuSCs, and followed subjects as they aged (Fig. 1B). Tamoxifen-treated *Cox10* (Pax7-Cre^ERT2^; Co×10^flox/flox^) animals had a median lifespan of 578 days, and a maximum lifespan of 805 days, which was significantly shorter than tamoxifen-treated wild-type (wt) animals (Pax7-Cre^ERT2^; Co×10^+/+^, median 861.5 days, maximum 965 days) (Fig. 1B). Histologic examination at 12 months of age did not reveal signs of inflammation or fibrosis in muscles from *Cox10* animals (Fig. S1C,D). However, *Cox10* animals exhibited decreased size of the tibialis anterior (TA) muscle and decreased cross-sectional area of myofibers in multiple muscle groups (Fig. 1C-E, Fig. S1E). Myofibers from *Cox10* animals had significant impairments in mitochondrial cytochrome c oxidase (COX) activity (Fig. 1F, Fig. S1F). *Cox10* animals also exhibited deficits in muscle performance, including decreased grip-strength and maximal running distance (Fig. 1G,H). Thus, inducing ETC dysfunction in MuSCs is sufficient to decrease muscle and myofiber size, motor performance and lifespan as mice age.

**Figure 1:**
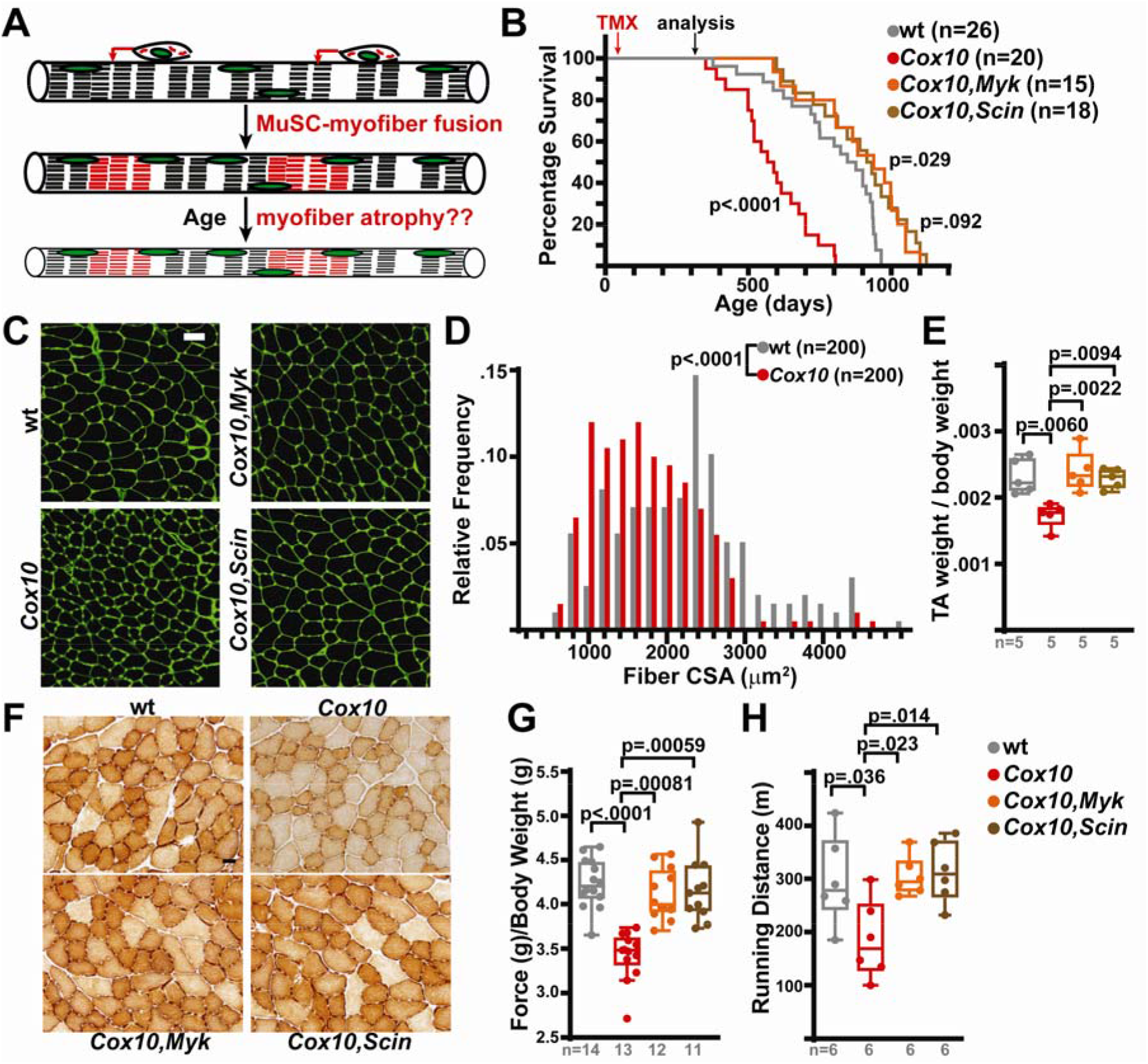
Complex IV dysfunction in MuSCs induces pre-mature sarcopenia and limits lifespan in mice. A) Overall model: Induced mitochondrial dysfunction in MuSCs triggers them to fuse into adjacent adult myofibers (red arrows) which initiates sarcopenic features. B) Survival curve of mice with the indicated genotype, following tamoxifen administration at 6 weeks of age. A subset of mice were euthanized at 12 months of age for analysis. p-values represent comparison with wt survival curves. C) Representative immunofluorescent images from tibialis anterior (TA) cross-sections of animals at 12 months of age. Muscle boundaries are visualized by staining with WGA (green). Scale bar, 50 μm. D) Histogram of myofiber cross-sectional area from TA muscles of wild-type and *Cox10* animals. E) TA muscle weight (normalized to body weight) for animals of the indicated genotype at 12 months of age. F) Representative histochemistry of COX activity in muscle cross-sections from TA muscles at 12 months of age. Scale bar, 50 μm. G) Hindlimb grip strength measurements from animals at 12 months of age. H) Maximal treadmill distance of animals at 12 months of age. Box plots indicate the inter-quartile ranges; whiskers are plotted with the Tukey method. Statistical analysis was performed using 1-way ANOVA (E,H), Kruskal-Wallis (G), log-rank test (B), or Kolmogorov-Smirnov test (D), with adjustments for multiple comparisons.

In parallel, we assessed animals in which Cox10 was co-deleted with either Myomaker or Scinderin in the MuSC compartment (‘*Cox10,Myk’* or ‘*Cox10,Scin’*). Myomaker is a transmembrane protein required for MuSC fusion, while Scinderin is an actin-network re-organizing protein that is specifically required for MuSC-myofiber fusion (*5, 13, 14*). *Cox10,Myk* and *Cox10,Scin* animals did not experience shortened lifespans, myofiber or muscle atrophy, mitochondrial dysfunction, or motor function deficits (Fig. 1B-H), indicating that MuSC-myofiber fusion is required for the initiation of sarcopenic-features in this genetic model.

We observed the presence of COX-negative, SDH-positive (COX-SDH+) fibers in 12 month old *Cox10* muscle, similar to histologic characteristics observed in muscle biopsies of elderly human patients (*15-17*) (Fig. S1G). COX-SDH+ fibers were not observed in wild-type, *Cox10,Myk* or *Cox10,Scin* animals (Fig. S1H). These results indicate that loss of ETC function in MuSCs is sufficient to accelerate myofiber atrophy with age-associated histologic features, and suggest that inhibition of MuSC-myofiber fusion may be protective in age-associated atrophy.

We next assessed the role of MuSC-myofiber fusion during normal aging by comparing wild-type mice with animals where Myomaker or Scinderin were conditionally deleted in MuSCs at 6 weeks of age (*Myk:* Pax7-Cre^ERT2^; Myk^flox/flox^ or *Scin*: Pax7-Cre^ERT2^; Scin^flox/flox^). *Myk* and *Scin* animals exhibited extensions in maximal lifespan (1156 days (*Myk*) and 1178 days (*Scin*) vs. 965 days (wild-type)), without significant changes in median survival (Fig. 2A). In wild-type animals, myofibers maintain size between 2 and 12 months of age, followed by atrophy (Figure 2B,C). Myofiber atrophy was inhibited in muscles from *Scin* and *Myk* animals (Fig. 2B,C). Similarly, the absolute TA muscle weight increases from 2 to 12 months, but then declines by 24 and 30 months of age in wild-type animals (Fig. 2D). In *Scin* and *Myk* animals, muscle weight is retained at 24 and 30 months of age (Fig. 2D). We observed similar trends after normalizing for body weight (Fig. S1I).

**Figure 2:**
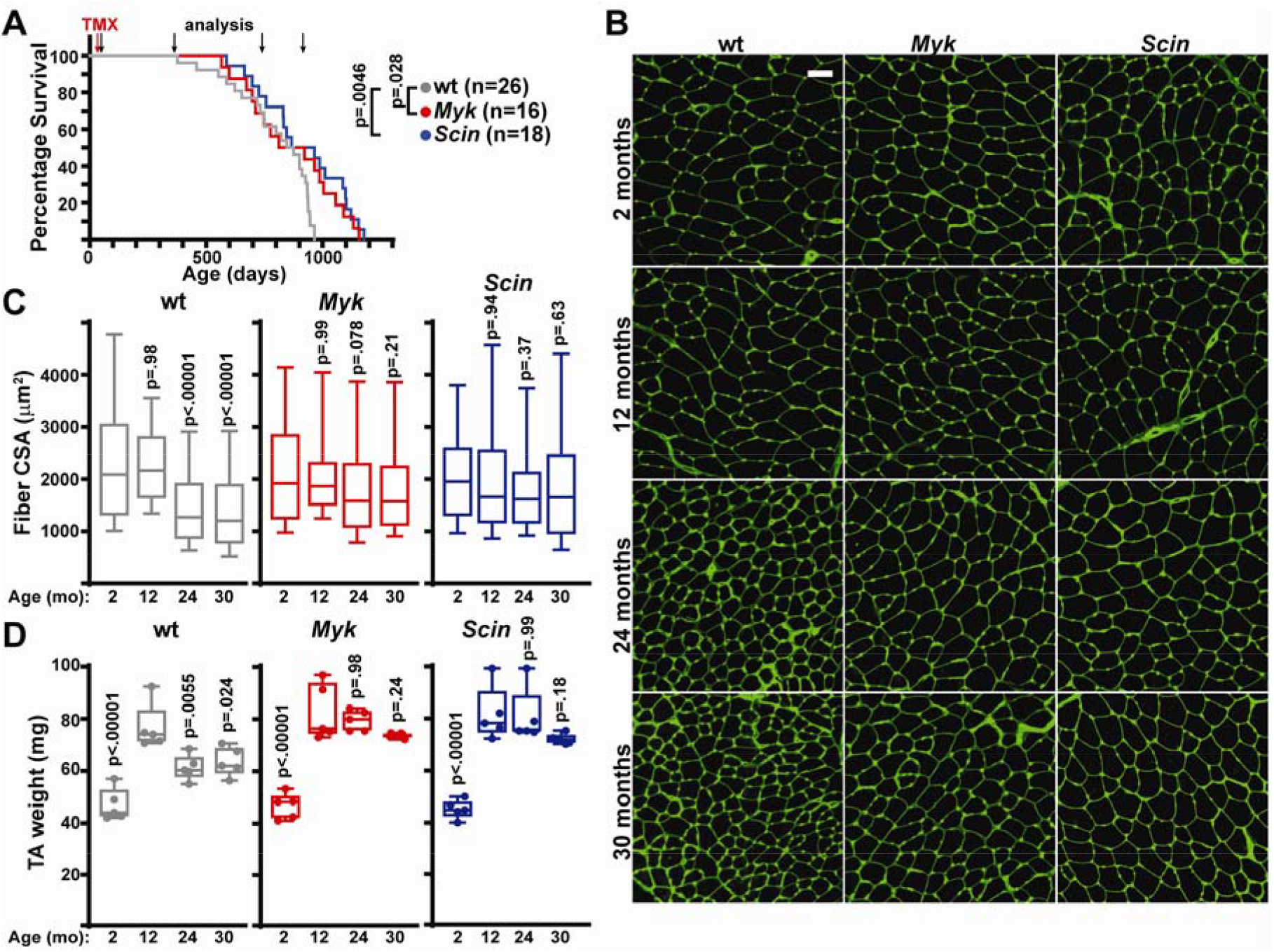
Inhibition of MuSC-myofiber fusion protects animals from myofiber atrophy during aging. A) Survival curves of mice of the indicated genotype, following tamoxifen administration at 6 weeks of age. p-values reflect comparisons with wild-type survival curve. A subset of animals were euthanized at 2, 12, 24 and 30 months of age for analysis. B) Representative immunofluorescent images from TA cross-sections. Muscle boundaries are visualized by WGA staining (green). Scale bar, 50 μm. C) Quantitation of myofiber cross-sectional area from TA muscles. 200 fibers were analyzed from n=5 mice per group. p-values reflect comparisons with 2 month old group. D) TA muscle weight at the indicated age. n=5 mice per group. p-values reflect comparisons with 12 month old group. Box plots indicate the inter-quartile ranges; whiskers indicate the 10^th^ and 90^th^ percentile. Statistical analysis was performed using 2-way ANOVA (C,D) or log-rank test (A) with adjustments for multiple comparisons.

Together, these results suggest a new model for the initiation and progression of myofiber atrophy during aging (Fig. 1A). MuSCs are known to accumulate mitochondrial genome mutations and dysfunction and are induced to fuse with neighboring myofibers with age (*5, 18, 19*). These fusion events clear damaged MuSCs from the stem cell population; however, adult myofibers accumulate the contents of damaged MuSCs, which accelerates the development of sarcopenic features and limits their maximal lifespan. Inhibiting MuSC-myofiber fusion blocks myofiber atrophy and can increase the maximal lifespan of mice. Interestingly, sporadic mitochondrial dysfunction within myofibers are a key feature of human sarcopenia (*15-17*). A number of studies indicate that segments of aged myofibers contain clonal expansions of mutant mitochondrial genomes that co-localize with electron transport chain defects (*20-22*), suggesting that mitochondrial defects within myofibers drive atrophy during aging (*23-25*). Thus, we propose that it is not the decline of MuSCs that significantly contributes to muscle aging, but instead the absorption of damaged mitochondrial genomes from aging MuSCs. Future work examining the biology and genotypes characteristic of aged MuSCs in human specimens will be important to validate this model.

## Acknowledgements

We thank Eric Olson for providing *Myk*^flox^ mice, and Michael Glogauer for providing *Scin*^flox^ mice.

## Author Contributions

X.W. and P.M. conceived the project. X.W. performed experiments. X.W. and P.M. prepared the figures and manuscript.

## Data availability

All data and materials are provided within the manuscript and supplementary materials, or upon request.

