## Supplemental Materials for "Fusion of dysfunction muscle stem cells with myofibers induces sarcopenia in mice"

**Experimental Procedures**

**Supplementary Figure S1**

### Experimental Procedures

#### Mice

Throughout this study, all genotypes refer to animals with conditional (flox) alleles, targeted to the muscle stem cell (MuSC) population. Animals with Cox10$^{flox}$ (strain 024697), Pax7-Cre$^{ERT2}$ (strain 017763), mito-Dendra2$^{flox}$ (strain 018385) alleles were purchased from the Jackson Laboratory. Animals with *Myk*$^{flox}$ and *Scin*$^{flox}$ alleles were kind gifts from Dr. Eric Olson and Dr. Michael Glogauer, and previously described (*12, 23*). All mice were maintained on C57BL6 backgrounds. The following genotypes are referred to in the manuscript:

| | |
|---|---|
| wt | Pax7-Cre$^{ERT2}$/Pax7-Cre$^{ERT2}$; mito-Dendra2$^{flox/flox}$ |
| *Cox10* | Pax7-Cre$^{ERT2}$/Pax7-Cre$^{ERT2}$; Cox10$^{flox/flox}$; mito-Dendra2$^{flox/flox}$ |
| *Cox10, Myk* | Pax7-Cre$^{ERT2}$/Pax7-Cre$^{ERT2}$; Cox10$^{flox/flox}$; Myk$^{flox/flox}$; mito-Dendra2$^{flox/flox}$ |
| *Cox10, Scin* | Pax7-Cre$^{ERT2}$/Pax7-Cre$^{ERT2}$; Cox10$^{flox/flox}$; Scin$^{flox/flox}$; mito-Dendra2$^{flox/flox}$ |
| *Myk* | Pax7-Cre$^{ERT2}$/Pax7-Cre$^{ERT2}$; Myk$^{flox/flox}$; mito-Dendra2$^{flox/flox}$ |
| *Scin* | Pax7-Cre$^{ERT2}$/Pax7-Cre$^{ERT2}$; Scin$^{flox/flox}$; mito-Dendra2$^{flox/flox}$ |

Both male and female mice were used in all experiments; if sex specific differences were not present, male and female mice were analyzed together. A random subset of animals were euthanized at the indicated timepoints for analysis; these animals were not included in the survival analysis. All mice were housed in the Animal Resource Center at the University of Texas Southwestern Medical Center under a 12 hr light-dark cycle and were fed ad libitum. All animal protocols were approved by the University of Texas Southwestern Institutional Animal Care and Use Committee.

For mouse injections, tamoxifen (TMX, Cayman Chemical, #132585) was dissolved in corn oil (Sigma-Aldrich, #C8267). 75mg/kg body mass was administered by intraperitoneal injection to 6-8-week-old mice, once per day for 5 consecutive days starting at 6 weeks of age.

Mice were euthanized and tissue harvested at various timepoints after tamoxifen administration as indicated.

For treadmill exercise capacity experiments, mice were acclimated to the treadmill (Columbus Instruments) by running 30 min/day for 3 days at slow speeds (up to 10m/min). On the day after acclimation, mice were run to exhaustion using the following protocol: The treadmill speed was initially set to 0 cm/sec and incremented by 5cm/sec, every 3 minutes until exhaustion. Exhaustion was determined by staying on the electrode more than 10 s as described previously (*24*), and the total running distance was recorded.

Mice hindlimb grip strength was measured using the following protocol (http://www.treat-nmd.eu/downloads/file/sops/dmd/MDX/DMD_M.2.2.001.pdf) by blinded laboratory staff. Briefly, mice were lifted and moved towards the bar of the force meter (Chatillon, #E-DFE-002) until they tightly gripped the bar with their hindlimbs. Mice were gently pulled away from the bar until the grasp was broken and the peak force measurement was recorded. 5 trials were repeated per mouse, and the high and low values were discarded. The average of the remaining force measurements was normalized to body weight.

Analysis of skeletal muscle

Following carbon dioxide asphyxiation and cervical dislocation, skeletal muscles were dissected, freshly frozen in liquid nitrogen cooled 2-methylbutane, and then embedded into O.C.T compound (Fisher Scientific, # 23-730-571). 10 μM sections were cut on a cryostat (Leica CM3050S). For H&E staining, slides were prepared following the protocol from the TREAT-NMD website (http://www.treat-nmd.eu/downloads/file/sops/cmd/MDC1A_M.1.2.004.pdf). For COX/SDH staining, slides were stained for COX activity and SDH activity as previously described (*25*). Sirius Red staining was performed following manufacturer's instructions (NovaUltra Picro-Sirius Red Stain Kit, #IW-3012). Sections were imaged using an Olympus IX83 microscope and analyzed with ImageJ software (NIH) to calculating COX intensity.

For immunofluorsence analysis, freshly frozen 10 μm cross-sections were fixed in formalin at room temperature for 5 min and stained with AlexaFluor 488 WGA (ThermoFisher, #W11261, 5 μg/mL) to visualize myofiber borders. Cross-sections were imaged on a Zeiss LSM780 inverted confocal microscope and analyzed with ImageJ software (NIH) to calculate myofiber cross-sectional area.

MuSC-myofiber fusion was assessed as previously described (*5, 26*), using lineage tracing with the mito-Dendra2 conditional allele. Skeletal muscle was fixed in formalin for 4 hours at room temperature, followed by overnight at 4°C. Muscles were rinsed with PBS and then mounted in a glass bottom dish (MatTek, # P35G-1.5-14-C). Muscles were imaged using a Zeiss LSM780 Inverted confocal microscope.

<u>Statistical analyses</u>

All data represent independent measurements from biological replicates. No statistical tests were used to predetermine sample size. Data sets for each group of measurement was tested for normality using the Shapiro-Wilk test. If the data was not normally distributed, the data was log-transformed and retested for normality. For normally-distributed data, groups were compared using the two-tailed Student's t-test (for 2 groups), or one-way ANOVA or two-way ANOVA (> 2 groups), followed by Tukey's or Dunnett's test for multiple comparisons. For data that was not normally distributed, we used non-parametric testing (Mann-Whitney or Kolmogorov-Smirnov tests for two groups and Kruskal-Wallis test for multiple groups), followed by Dunn's multiple comparisons adjustment. Multiple independent experiments with biological replicates were performed for all reported data, and the number of biological replicates are indicated in the figures.

**Figure S1**

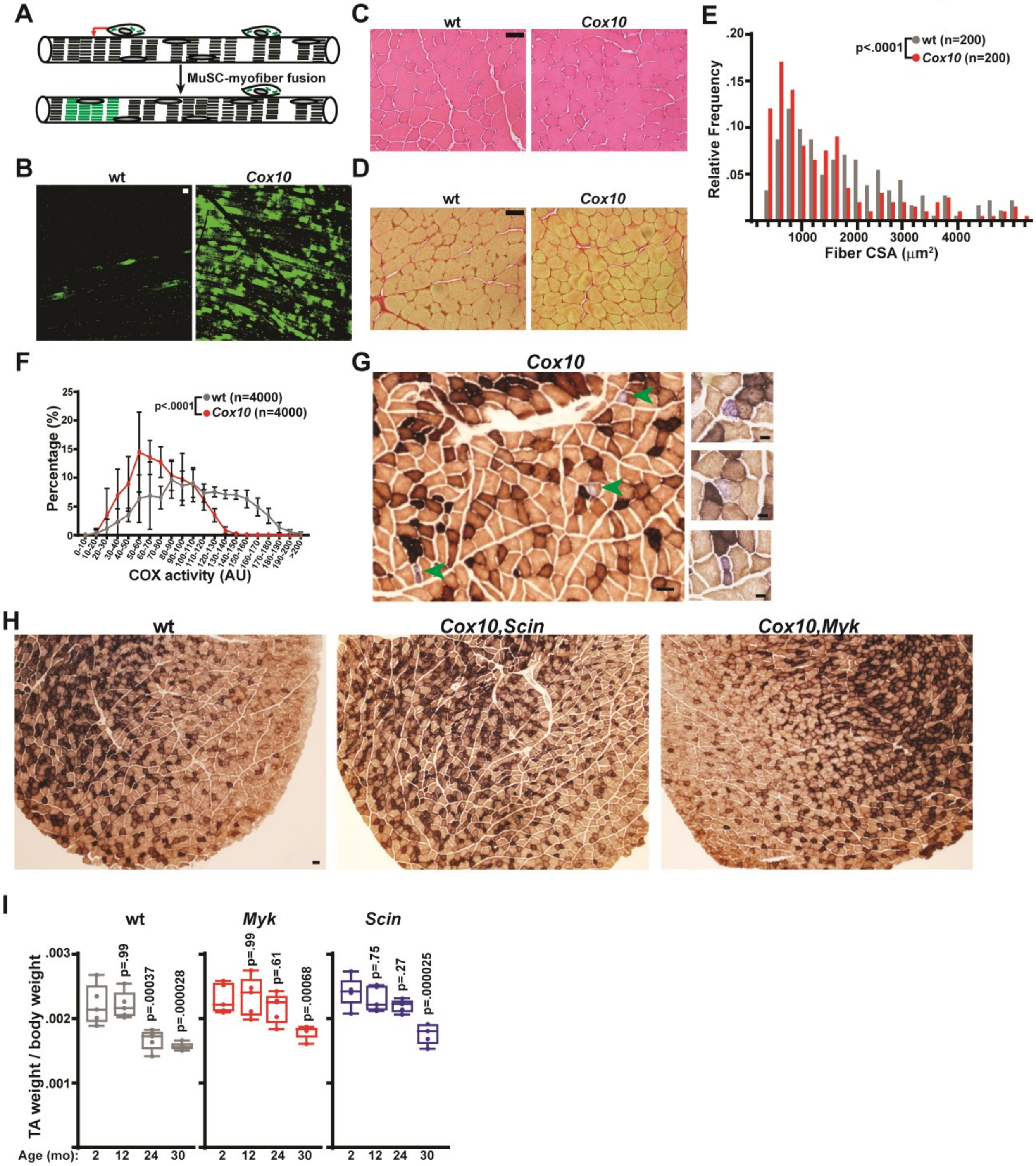


**Supplementary Figure S1: Complex IV dysfunction in MuSCs induces MuSC-myofiber fusion.**

A) Schematic for monitoring MuSC-myofiber fusion. MuSCs are lineage-traced using the mito-Dendra2 conditional allele combined with a Pax7-Cre$^{ERT2}$ allele (see Methods); this combination labels the mitochondria of MuSCs with Dendra2, which fluoresces in the green spectrum. As previously shown (*26*), if a MuSC-myofiber fusion event occurs (red arrow), mitochondria surrounding the fusion site are labeled with mito-Dendra2, which can be visualized on longitudinal sections. B) Representative longitudinal images of mito-Dendra2 labeling (green) in 12 month old wild-type and *Cox10* muscles. Scale bar, 50 μm. C) Representative H&E images of muscle cross-sections from tibialis anterior muscles in 12 month old animals. Scale bar, 50 μm. D) Representative images of muscle cross-sections from tibialis anterior muscles stained with Sirius Red to visualize collagen deposition. Scale bar, 50 μm. E) Histogram of myofiber cross-sectional area from gastrocnemius muscles of wild-type and *Cox10* animals at 12 months of age. F) Histogram of individual myofiber COX activity from TA muscles of the indicated genotype. 1000 fibers were analyzed per individual; n=4 mice per group. AU, arbitrary units. G) Representative low-magnification (left) and high-magnification (right) images of SDH-positive fibers (purple stain; green arrowheads) from 12month old *Cox10* animals. Scale bar, 20 µm. H) Representative immunohistochemistry of complex IV (COX; brown) and complex II (SDH; purple) activities in TA cross-sections from mice of the indicated genotype at 12 months of age. Scale bar, 50 µm. I) Tibialis anterior muscle weight (normalized to body weight) for animals of the indicated genotype and age. Statistical significance was assessed using Kolmogorov-Smirnov (E,F) or 2-way ANOVA (I) tests.
